# INKA, an integrative data analysis pipeline for phosphoproteomic inference of active phosphokinases

**DOI:** 10.1101/259192

**Authors:** Thang V. Pham, Robin Beekhof, Carolien van Alphen, Jaco C. Knol, Alex A. Henneman, Frank Rolfs, Mariette Labots, Evan Henneberry^1^, Tessa Y.S. Le Large, Richard R. de Haas, Sander R. Piersma, Henk M.W. Verheul, Connie R. Jimenez

## Abstract

Identifying (hyper)active kinases in cancer patient tumors is crucial to enable individualized treatment with specific inhibitors. Conceptually, kinase activity can be gleaned from global protein phosphorylation profiles obtained with mass spectrometry-based phosphoproteomics. A major challenge is to relate such profiles to specific kinases to identify (hyper)active kinases that may fuel growth/progression of individual tumors. Approaches have hitherto focused on phosphorylation of either kinases or their substrates. Here, we combine kinase-centric and substrate-centric information in an Integrative Inferred Kinase Activity (INKA) analysis. INKA utilizes label-free quantification of phosphopeptides derived from kinases, kinase activation loops, kinase substrates deduced from prior experimental knowledge, and kinase substrates predicted from sequence motifs, yielding a single score. This multipronged, stringent analysis enables ranking of kinase activity and visualization of kinase-substrate relation networks in a biological sample. As a proof of concept, INKA scoring of phosphoproteomic data for different oncogene-driven cancer cell lines inferred top activity of implicated driver kinases, and relevant quantitative changes upon perturbation. These analyses show the ability of INKA scoring to identify (hyper)active kinases, with potential clinical significance.

## Introduction

Cancer is associated with aberrant kinase activity (Hanahan & Weinberg, 2011), and among recurrently altered genes some 75 encode kinases that may “drive” tumorigenesis and/or progression (Vogelstein *et al*, 2013). In the last decade, multiple kinase-targeted drugs including small-molecule inhibitors and antibodies have been approved for clinical use in cancer treatment (Knight *et al*, 2010). However, even when selected on the basis of extensive genomic knowledge, only a subpopulation of patients experiences clinical benefit (Huang *et al*, 2014; Valabrega *et al*, 2007; Flaherty *et al*, 2010), while invariably resistance also develops in responders. Resistance cannot only result from mutations in the targeted kinase or downstream pathways, but also from alterations in more distal pathways (Al-Lazikani *et al*, 2012; Trusolino & Bertotti, 2012; Ramos & Bentires-Alj, 2015). This complexity calls for tailored therapy based on detailed knowledge of the individual tumor’s biology, including a comprehensive profile of (hyper)active kinases. MS-based phosphoproteomics enables global protein phosphorylation profiling of cells and tissues (Jimenez & Verheul, 2014; Casado *et al*, 2016), but to arrive at a prioritized list of actionable (combinations of) active kinases, a dedicated analysis pipeline is required as the data are massive and complex. Importantly, a prime prerequisite for personalized treatment requires that the analysis is based on a single sample.

Different kinase ranking approaches have been described previously. Rikova and colleagues sorted kinases on the basis of the sum of the spectral counts (an MS correlate of abundance) for all phosphopeptides attributed to a given kinase, and identified known and novel oncogenic kinases in lung cancer (Rikova *et al*, 2007). This type of analysis can be performed in individual samples, but is limited by a focus on phosphorylation of the kinase itself, rather than the (usually extensive) set of its substrates. Instead, several substrate-centric approaches also exist, including KSEA (Casado *et al*, 2013; Terfve *et al*, 2015; Wilkes *et al*, 2015), pCHIPS (Drake *et al*, 2016), and IKAP (Mischnik *et al*, 2016).

Neither a kinase-centric nor a substrate-centric phosphorylation analysis may suffice by itself to optimally single out major activated (driver) kinase (s) of cancer cells. To achieve an optimized ranking of inferred kinase activities based on MS-derived phosphoproteomics data for single samples, we propose a multipronged, rather than a singular approach. In this study, we devised a phosphoproteomics analysis tool for prioritizing active kinases in single samples, called Integrative Inferred Kinase Activity (INKA) scoring. INKA combines direct observations on phosphokinases (either all kinase-derived phosphopeptides or activation loop peptides specifically), with observations on phosphoproteins that are known or predicted substrates for the pertinent kinase. To demonstrate its utility, we analyzed cancer cell line ‘cases’ with known driver kinases in a single-sample manner, such as would be required in a personalized medicine setting and explored dynamic changes upon perturbation.

## Results

### INKA: integration of kinase-centric and substrate-centric evidence to infer kinase activity from single-sample phosphoproteomics data

To infer kinase activity from phosphoproteomics data of single samples, we developed a multipronged data analysis approach. Figure 1 summarizes the data collection (Fig 1A) and analysis workflows (Fig 1B) of the current study. For in-house data generation, we utilized phosphotyrosine (pTyr)-based phosphoproteomics of cancer cell lines and tumor needle biopsies (Supplementary Table 1). Kinases covered by individual analysis approaches are detailed in Supplementary Table 2.

**Figure 1.**
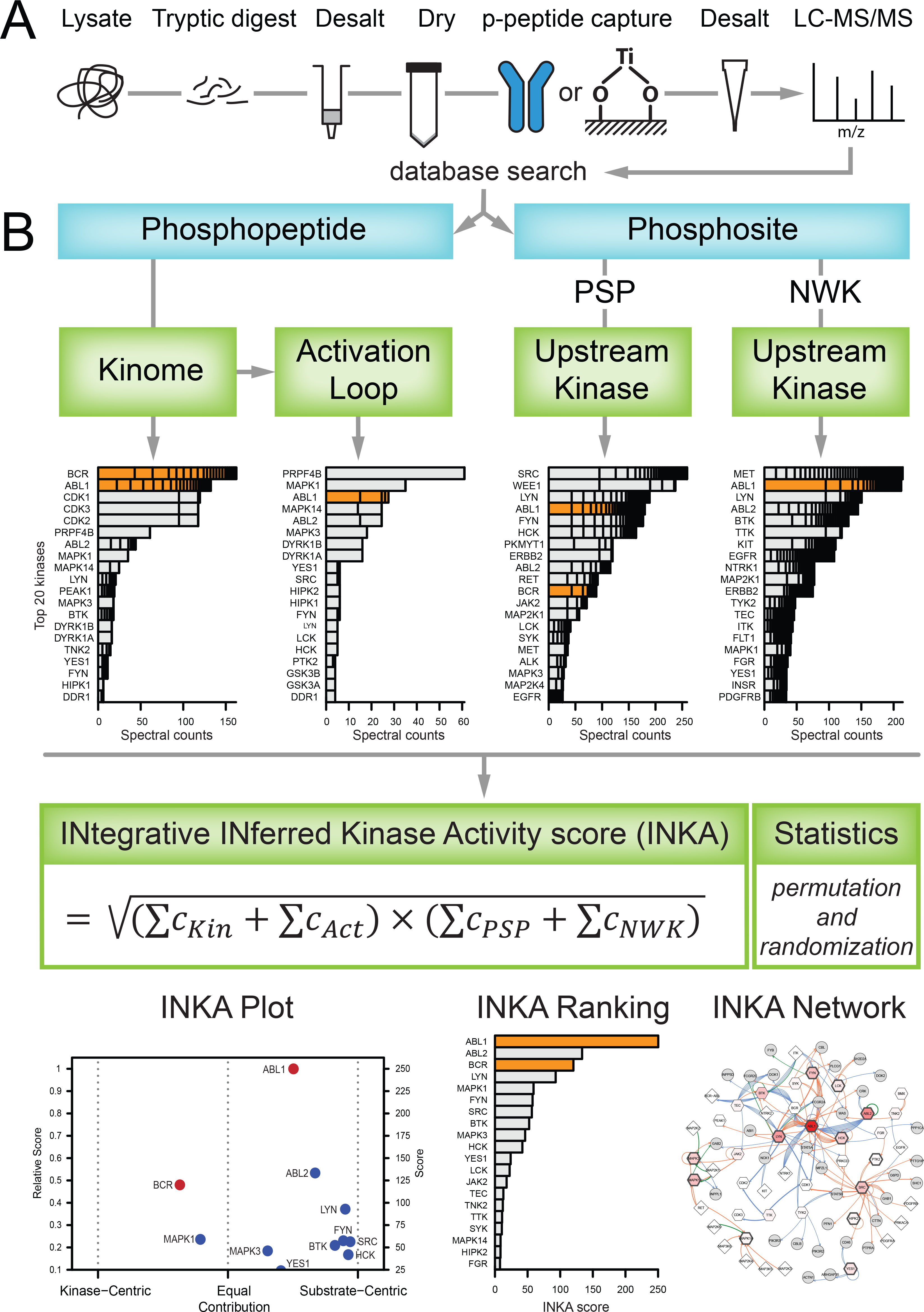
Generic phosphoproteomics workflow and data analysis strategy. A Overview of an MS-based phosphoproteomics experiment. Proteins from a biological sample are digested with trypsin, and phosphopeptides are enriched for analysis by (orbitrap-based) LC-MS/MS. Phosphopeptides can be captured with various affinity resins; here, data were analyzed of phosphopeptides enriched with anti-phosphotyrosine antibodies and TiOx. Database-based phosphopeptide identification, and phosphosite localization and quantification is performed using a tool like MaxQuant. B Scheme of INKA analysis for identification of active kinases in a single biological sample. Quantitative phosphodata for established kinases are taken as direct (kinase-centric) evidence, using either all phosphopeptides attributed to a given kinase (’Kinome’), or only those from the kinase activation loop segment (’Activation Loop’). Phosphosites are filtered for class I phosphosites (localization probability > 75%) (Olsen *et al*, 2006), coupled to phosphopeptide spectral count data, and used for substrate-centric inference of kinases on the basis of kinase-substrate relationships that are either experimentally observed (provided by PhosphoSitePlus, ‘PSP’) or predicted by an algorithm using sequence motif and protein-protein network information (NetworKIN, ‘NWK’). All evidence lines are integrated in a kinase-specific INKA score using the geometric mean of combined spectral count data (’C’) for kinase-centric and substrate-centric modalities. Results are visualized in a scatter plot of INKA scores for kinases scoring ≥10% of the maximum (’INKA Plot ‘; horizontal shifts from the middle indicate evidence being more kinase-centric or more substrate-centric). For top 20 INKA-scoring kinases, a score bar graph (’INKA Ranking’), and a kinase-substrate relation network for pertinent kinases and their observed substrates (’INKA Network’) are also produced.

First, phosphopeptides derived from established protein kinases (KinBase, http://kinase.com) (Manning *et al*, 2002) were analyzed. Kinase hyperphosphorylation is commonly associated with increased kinase activity. This is the rationale for using the sum of spectral counts (the number of identified MS/MS spectra) for all phosphopeptides derived from a kinase as a proxy for its activity, and to rank kinases accordingly, as pioneered by Rikova *et al.* (Rikova *et al*, 2007)

Second, kinase activation loop phosphorylation was analyzed. Most kinases harbor an activation segment, residing between highly conserved Asp-Phe-Gly and Ala-Pro-Glu motifs. Phosphorylation of residues in the activation loop counteracts the positive charge of a critical arginine in the catalytic loop, eliciting conformational changes and consequent kinase activation (Nolen *et al*, 2004). To identify phosphopeptides that are derived from a kinase activation segment, we used the Phomics toolbox (http://phomics.jensenlab.org) (Munk *et al*, 2016). Subsequently, kinases were ranked after spectral count aggregation as described above.

Third, as a substrate-centric complement to the kinase-centric analyses above, and similar to a key ingredient in KSEA analysis (Casado *et al*, 2013), one can backtrack phosphorylation of substrates to responsible kinases as an indirect way to monitor kinase activity. Experimentally established kinase-substrate relationships listed by PhosphoSitePlus (Hornbeck *et al*, 2015) were used to link substrate-associated spectral counts to specific kinases, followed by kinase ranking.

Fourth, another substrate-centric analysis was included to complement the previous step. To date, databases logging experimental kinase-substrate relationships are far from complete, leaving a large proportion of phosphopeptides that cannot be mapped as a kinase substrate through that avenue. Therefore, we applied the NetworKIN prediction algorithm (Linding *et al,* 2007; Horn *et al*, 2014) to observed phosphosites to generate a wider scope of kinase-substrate relationships. NetworKIN uses phosphorylation sequence motifs and protein-protein network (path length) information to predict and rank kinases that may be responsible for phosphorylation of specific substrate phosphosites. After applying score cutoffs to restrict the NetworKIN output to the most likely kinase-substrate pairs, kinases were ranked by the sum of all spectral counts associated with their predicted substrates.

Finally, we devised a method to integrate the four analyses as described above and to provide a single metric for pinpointing active kinases in biological samples analyzed by phosphoproteomics (Fig 1B, Materials and Methods). For a given kinase, associated values in either of the two kinase-centric analyses are summed, and the same is done for the two substrate-centric analyses. Subsequently, the geometric mean of both sums is taken as an integrated inferred kinase activity, or INKA score. A non-zero INKA score requires both kinase-centric and substrate-centric evidence to be present. Furthermore, a skew parameter is calculated (0 for exclusively kinase-centric, 1 for exclusively substrate-centric, and 0.5 for equal contribution; see Materials and Methods) indicating to which extent the INKA score is derived from kinase-centric or from substrate-centric evidence. For kinases that are missing from PhosphoSitePlus and can not be inferred by NetworKIN prediction as well, a separate kinase-centric ranking is performed to include these MS-observed enzymes in the analysis. The latter involves 172 out of 538 established protein kinases considered in our analyses (Supplementary Fig 1 and Supplementary Table 2). For kinases inferred through PhosphoSitePlus/NetworKIN but not observed by MS, the reciprocal analysis is not performed as kinases display overlapping substrate specificities precluding unequivocal assignment of a substrate to a specific kinase.

Results are visualized through different plots (Fig 1B). Individual analyses each result in a bar graph with top 20 kinases. Integration by INKA scoring results in a scatter plot for all kinases with an INKA score that is at least 10% of the top scoring kinase (vertical: INKA score, horizontal: skew towards kinase-centric or substrate-centric evidence). For the top 20 kinases (by INKA score), a ranked bar graph and a network of all inferred kinase-substrate connections are visualized as well (Fig 1B). This pipeline is available to users who can upload their data to www.inkascore.org for INKA scoring.

### INKA analysis of cancer cell line use cases

To assess performance of the INKA approach, we carried out pTyr-based phosphoproteomics with four well-studied cell lines for which oncogenic driver kinases are known: K562 chronic myeloid leukemia (CML) cells harboring a *BCR-ABL* fusion (Klein *et al*, 1976; Heisterkamp *et al*), SK-Mel-28 melanoma cells carrying mutant *BRAF* (Carey *et al*, 1976; Davies *et al*, 2002), HCC827-ER3 lung carcinoma cells with mutant *EGFR* (Girard *et al*, 2000; Zhang *et al*, 2012), and H2228 lung carcinoma cells with an *EML4-ALK* fusion (Soda *et al*, 2007; Rikova *et al,* 2007). Figure 2 displays, on a single row per cell line, bar graphs with the top 20 kinases for each of the four basic analyses (Kinome, Activation Loop, PhosphoSitePlus, and NetworKIN) as well as combined score analysis (INKA). Bars for known driver kinases are highlighted by coloring for each cell line except SK-Mel-28, which is driven by BRAF, a serine/threonine kinase that cannot be detected by pTyr-based phosphoproteomics. In the latter case, downstream driver targets in the MEK-ERK pathway (MAP2K1, MAP2K2, MAPK1, MAPK3) are highlighted (Fig 2B). Underlying data can be found in Supplementary Table 3.

**Figure 2.**
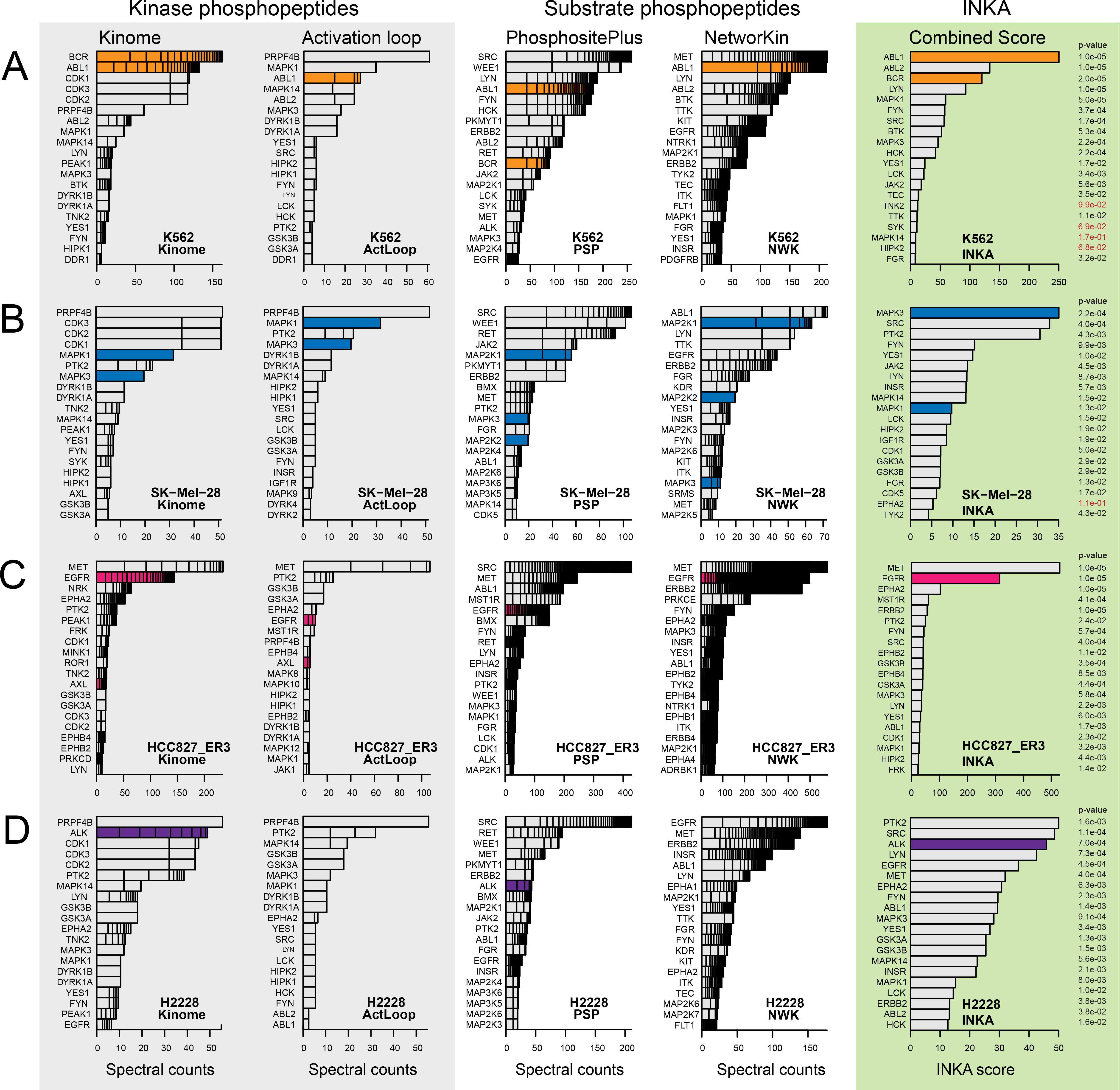
Ranking of top 20 kinases in four cell line use cases by each of four lines of evidence and integrative INKA scoring. A K562 chronic myelogenous leukemia cells with a *BCR-ABL* fusion. INKA score ranking indicates that ABL1/BCR-ABL (orange bars) exhibits principal kinase activity in this cell line, in line with a role as an oncogenic driver. B SK-Mel-28 melanoma cells with mutant *BRAF*. In the ‘Kinome’ analysis, CDK1, CDK2 and CDK3 share a second place, based on phosphopeptides that cannot be unequivocally assigned to either of them. INKA scoring implicates MAPK3 as the number one activated kinase. As SK-Mel-28 is driven by BRAF, a serine/threonine kinase that is missed by pTyr-based phosphoproteomics, downstream targets in the MEK-ERK pathway are highlighted by blue coloring. C Erlotinib-resistant HCC827-ER3 NSCLC cells with mutant *EGFR*. INKA scores reveal the driver EGFR (pink coloring) as second-highest ranking and MET as highest ranking kinase, respectively. D H2228 NSCLC cells with an *EML4-ALK* fusion. The driver ALK (purple coloring) is ranked as a top 3 kinase by INKA score, slightly below PTK2 and SRC. Data information: For each cell line, bar graphs depict kinase ranking based on kinase-centric analyses (panel “Kinase phosphopeptides”), substrate-centric analyses (panel “Substrate phosphopeptides”), and combined scores (panel “INKA”). Bar segments represent the number and contribution of individual phosphopeptides (kinase-centric analyses) or phosphosites (substrate-centric analyses). P-values flanking INKA score bars were derived through a randomization procedure with 10^5^ permutations of both peptide-spectral count links and kinase-substrate links.

From the first column of bar graphs in Fig 2 it is evident that kinome analysis, based on all phosphopeptides identified for a kinase, is a strong indicator for (hyper)active kinases, as was found previously (Rikova *et al*, 2007; Guo *et al*, 2008). All tyrosine kinase drivers rank among the top two. Activation loop analysis (Fig 2, second column) follows a similar trend, but suffers from a focus on a small protein segment. With respect to substrate-centric analyses, both PhosphoSitePlus and NetworKIN analyses (Fig 2, third and fourth columns, respectively) rank driver kinases among the top 10. One of the two analyses often outperforms the other, but not in a consistent pattern, underscoring our choice to use a combination of these approaches. Since substrate-centric inference attributes data from multiple, possibly numerous, substrate phosphosites to a single kinase, bar segments may coalesce into a black stack in more extreme cases.

From the above it is clear that individual analysis modules harbor complementary layers of information about phosphokinases and phosphosubstrates that can be integrated using INKA scoring into a more balanced and accurate pattern of inferred kinase activity (Fig 2, last column). Importantly, all driver (-related) kinases rank among the INKA top 3 for the tested cell lines (with MAPK3 serving as a surrogate for the BRAF Ser/Thr kinase in SK-Mel-28).

In order to explore the statistical significance of the INKA score, and especially the impact of kinase inference through substrate-centric PhosphoSitePlus and NetworKIN analyses, we permuted both phosphopeptide-spectral count links (for a sample) and kinase-substrate links (for all PhosphoSitePlus and NetworKIN data), followed by INKA score calculation. After 10^5^ permutations, sample- and kinase-specific INKA score distributions were obtained and used to derive p-values for each actual INKA score. Based on this calculation, almost all top 20 INKA scores were significant (Fig 2). Higher INKA scores clearly correlated with lower p-values (Supplementary Fig 2), underscoring the relevance of the INKA score.

### Linking INKA analysis to cell biology and treatment selection

To assess to what extent (hyper)active phosphokinases identified by the above INKA analysis represent actionable drug targets, we investigated public cell line drug sensitivity data from the ‘Genomics of Drug Sensitivity in Cancer’ resource (Yang *et al*, 2013) (GDSC, http://www.cancerrxgene.org, Supplementary Table 4).

#### K562

Clinically, treatment of BCR-ABL-positive CML patients with kinase inhibitors such as nilotinib, imatinib, and dasatinib has shown great benefit (Kantarjian *et al*, 2010; Saglio *et al,* 2010; Larson *et al*, 2012), in line with addiction of tumor cells to the BCR-ABL fusion kinase. BCR/ABL activity was inferred in all basic analyses of K562, causing ABL1 to stand out as a prime candidate in the INKA analysis (Fig 2A, Fig 3A). The integrated analysis also inferred phosphorylation of downstream signaling partners of ABL1 such as SRC-family kinases and MAPK1/3 among all members of the kinase-substrate relation network for top 20 INKA kinases in K562 (Fig 3A, enlarged in Supplementary Fig 3). As expected, GDSC data indicate K562 cells to be sensitive to various ABL inhibitors (Supplementary Table 4).

**Figure 3.**
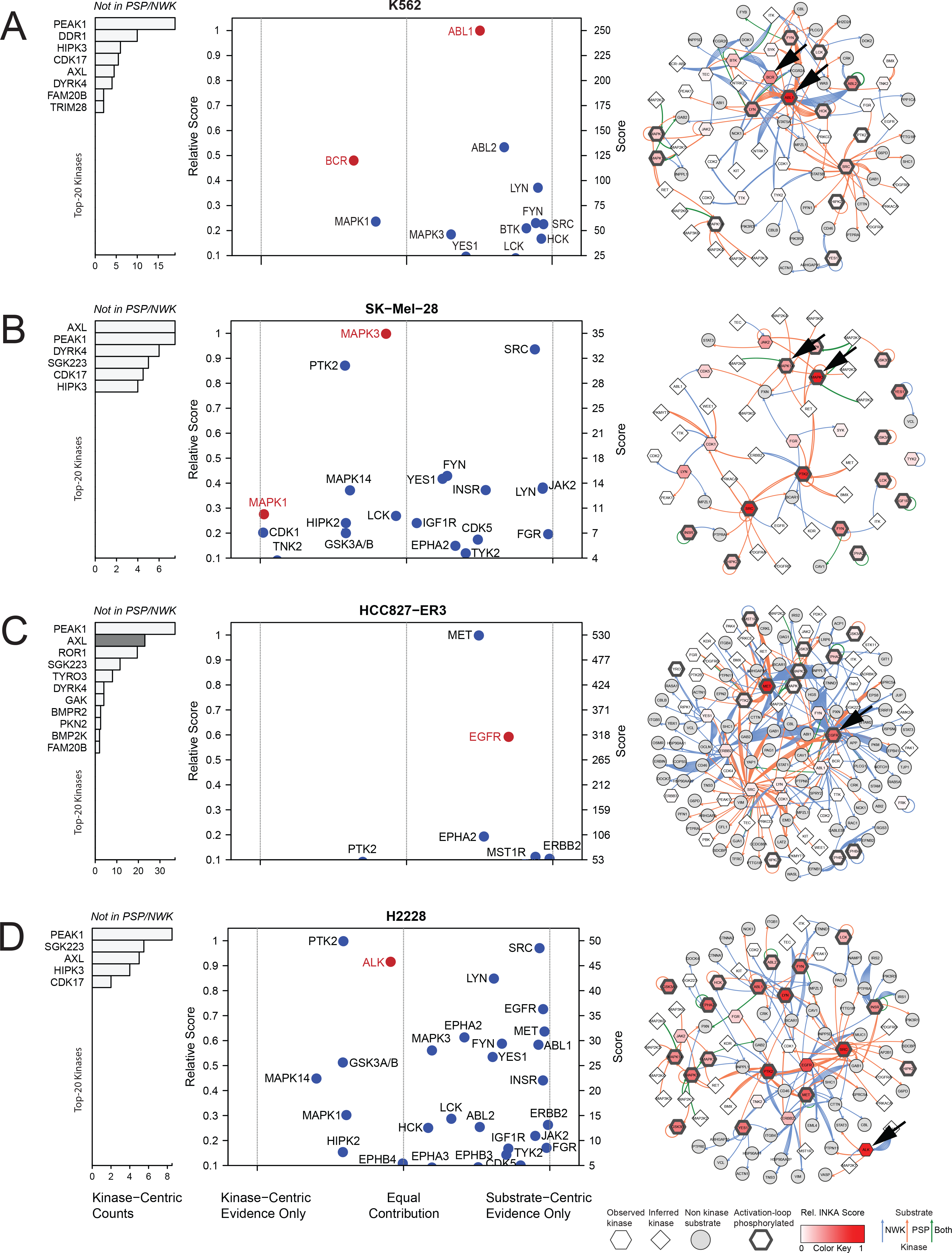
INKA plots and kinase-substrate relation networks for four cell line use cases. A K562 CML cells with a *BCR-ABL* fusion. ABL1 is the most activated kinase, with relatively equal contributions from both analysis arms. It is a highly connected, central node in the network. B SK-Mel-28 melanoma cells with mutant *BRAF*. Downstream MEK-ERK pathway members are highlighted in lieu of BRAF which is missed by the current pTyr-based workflow. MAPK3 is the top activated kinase. The network includes two clusters with highly connected activated kinases, MAPK1/3 and SRC, respectively. C Erlotinib-resistant HCC827-ER3 NSCLC cells with mutant *EGFR*. EGFR and MET are the most active, highly connected kinases. AXL, inactive in parental cells (see Supplementary Fig **6**), but associated with erlotinib resistance in this sub-line, can only be analyzed through the kinase-centric arm (pink bar highlighting). D H2228 NSCLC cells with an *EML4-ALK* fusion. ALK is a high-ranking kinase with roughly equal evidence from both analysis arms. Multiple highly active and connected nodes imply relative insensitivity to ALK inhibition, in line with previous functional data. Larger networks are shown in Supplementary Figs 3–5 and Supplementary Fig **7**. Data information: In INKA plots proper, the vertical position of kinases (drivers in red) is determined by their INKA score, whereas the horizontal position is determined by the (im)balance of evidence from kinase-centric and substrate-inferred arms of the analysis. Kinases not covered by PhosphoSitePlus (PSP) and NetworKIN (NWK) are visualized in a flanking bar graph.

#### SK-Mel-28

A vast majority of patients with *BRAF*-mutant melanoma carry a mutation encoding an activating V600E substitution in the BRAF kinase domain (Forbes *et al*, 2015; Lovly *et al,* 2012; Rubinstein *et al*, 2010). Treatment with small-molecule inhibitors of BRAF (Chapman *et al*, 2011; Hauschild *et al*, 2012; Flaherty *et al*, 2012) and/or downstream MEKs (MAP2K1/2) (Flaherty *et al*, 2012; Robert *et al*, 2015) often induces strong initial responses, but both single-agent and combination treatments are commonly hampered by development of resistance. The BRAF Ser/Thr kinase cannot be directly observed with pTyr-based phosphoproteomics, but INKA analysis of SK-Mel-28 revealed high- and medium-ranking activity of its downstream pathway members MAPK3 and MAPK1, respectively, in addition to high activity of focal adhesion kinase (PTK2) and SRC-family members (Fig 2B, Fig 3B). Of the latter, SRC is a central node in the inferred kinase-substrate network of top 20 INKA-scoring kinases in SK-Mel-28 (Fig 3B, Supplementary Fig 4). GDSC data indicate that inhibition of BRAF and downstream MEK1/2 is successful in reducing SK-Mel-28 growth, while inhibition more downstream of MAPK1/3 is less effective (Supplementary Table 4). Single-agent inhibition was not effective for PTK2, and undocumented for SRC-family kinases FYN and YES1. Based on INKA data, one could test ‘combination therapy’ with a PTK2 or SRC inhibitor and a BRAF or MEK1/2 inhibitor. Interestingly, a previous study showed the importance of SRC in BRAF inhibitor resistance in *BRAF*-mutant melanoma cells and patient-derived tissues (Girotti *et al*, 2013).

#### HCC827-ER3

Non-small cell lung carcinoma (NSCLC)-derived HCC827 cells expressing mutant EGFR^E746-A750^ are highly sensitive to EGFR inhibitors (GDSC, Supplementary Table 4). HCC827-ER3 is a sub-line of HCC827 with *in vivo* acquired resistance to the EGFR inhibitor erlotinib (Zhang *et al*, 2012), and cross-resistance to sunitinib (van der Mijn *et al*, 2016). INKA analysis suggests that HCC827-ER3 still contains hyperactive EGFR, with its score ranking second (Fig 2C, Fig 3C), indicating driver potential. Network visualization of inferred kinase-substrate relations (Fig 3C, Supplementary Fig 5) shows that both EGFR and MET are highly connected hyperactive nodes. Based on INKA scoring, acquired resistance of HCC827-ER3 to erlotinib (Zhang *et al*, 2012; van der Mijn *et al*, 2016) could potentially be overcome by co-targeting MET, which ranks first. However, high MET phosphorylation was already found in parental, erlotinib-sensitive HCC827 cells (Zhang *et al*, 2012; van der Mijn *et al*, 2016) (Supplementary Table 3E, Supplementary Fig 6), that appear relatively insensitive to MET inhibitors (Supplementary Table 4). Treating HCC827 with gefitinib, which inhibits EGFR but not MET (Davis *et al*, 2011), not only reduces hyperphosphorylation but also overall levels of MET (Guo *et al*, 2008). Thus, high INKA ranking of MET may be the result of EGFR-MET crosstalk, rather than indicating driver activity. Interestingly, growth of HCC827-ER3 is inhibited after co-targeting of EGFR and AXL (Guo *et al*, 2008; van der Mijn *et al*, 2016). Unfortunately, AXL (hyper)activity cannot be pinpointed by INKA, as both experimental (PhosphoSitePlus) and predicted (NetworKIN) kinase-substrate relationship data are lacking for AXL. However, in a kinase-centric ranking devised for kinases that suffer from such a problem, phospho-AXL ranks second with relatively high counts (Fig 2C, Fig 3C left panel) whereas it was not identified in parental HCC827 cells (Supplementary Fig 6), in line with AXL being a resistance hub in HCC827-ER3 (Guo *et al*, 2008; van der Mijn *et al*, 2016).

#### H2228

In NSCLC-derived H2228 cells, an *EML4-ALK* fusion underlies oncogenic ALK kinase activity. Indeed, several ALK inhibitors have been approved for the treatment of ALK fusion-positive NSCLC. ALK ranked third in INKA analysis (Fig 2D, Fig 3D), which may be due to paucity of information for this kinase (as for AXL, see above) or indicate involvement of other kinases in fueling cell growth. GDSC data indicate that effective inhibitors of H2228 growth are rare, even when targeting ALK (alectinib: IC50 4.4 µM, Z-score −1.92; Supplementary Table 4). The kinase-substrate network for top 20 INKA-scoring kinases in H2228 (Fig 3D, Supplementary Fig 7) shows that in addition to ALK there are multiple hyperactive, highly connected kinases (e.g., PTK2, SRC, EGFR) as candidates for combination treatment. Inhibitors targeting these kinases individually do not have a significant effect on H2228 proliferation (Supplementary Table 4). PTK2 and SRC are inhibited by several ALK inhibitors at IC50 concentrations (Davis *et al*, 2011), ruling these kinases out as co-targets. Interestingly, EGFR, a high-ranking kinase in INKA and a central node in the network, is implicated in reduced sensitivity to ALK inhibition in H2228, while dual inhibition of ALK and EGFR results in highly increased apoptosis (Voena *et al*, 2013). Furthermore, both simultaneous knockdown and combined drug inhibition of these kinases inhibit H2228 cell proliferation and tumor xenograft growth more potently than targeting either kinase alone (Li *et al*, 2016).

### Testing the INKA approach with literature data

To further test our strategy for prioritizing targeting candidates, we also examined phosphoproteome data from the literature (Guo *et al*, 2008; Bai *et al*, 2012) (Fig 4, Supplementary Table 3). INKA analysis of data on *EGFR*-mutant NSCLC cell line H3255 (Guo *et al*, 2008) uncovered major EGFR activity in these cells, with EGFR ranking first, followed by MET (Fig 4A). In another study, the rhabdomyosarcoma-derived cell line A204 was associated with PDGFRα signaling (Bai *et al*, 2012), and INKA scoring of the underlying data accordingly ranks PDGFRα in second place (Fig 4B). In the same study, osteosarcoma-derived MNNG/HOS cells were shown to be dependent on MET signaling and sensitive to MET inhibitors (Bai *et al*, 2012). In line with this, INKA analysis clearly pinpointed MET as the major driver candidate in this cell line (Fig 4C).

**Figure 4.**
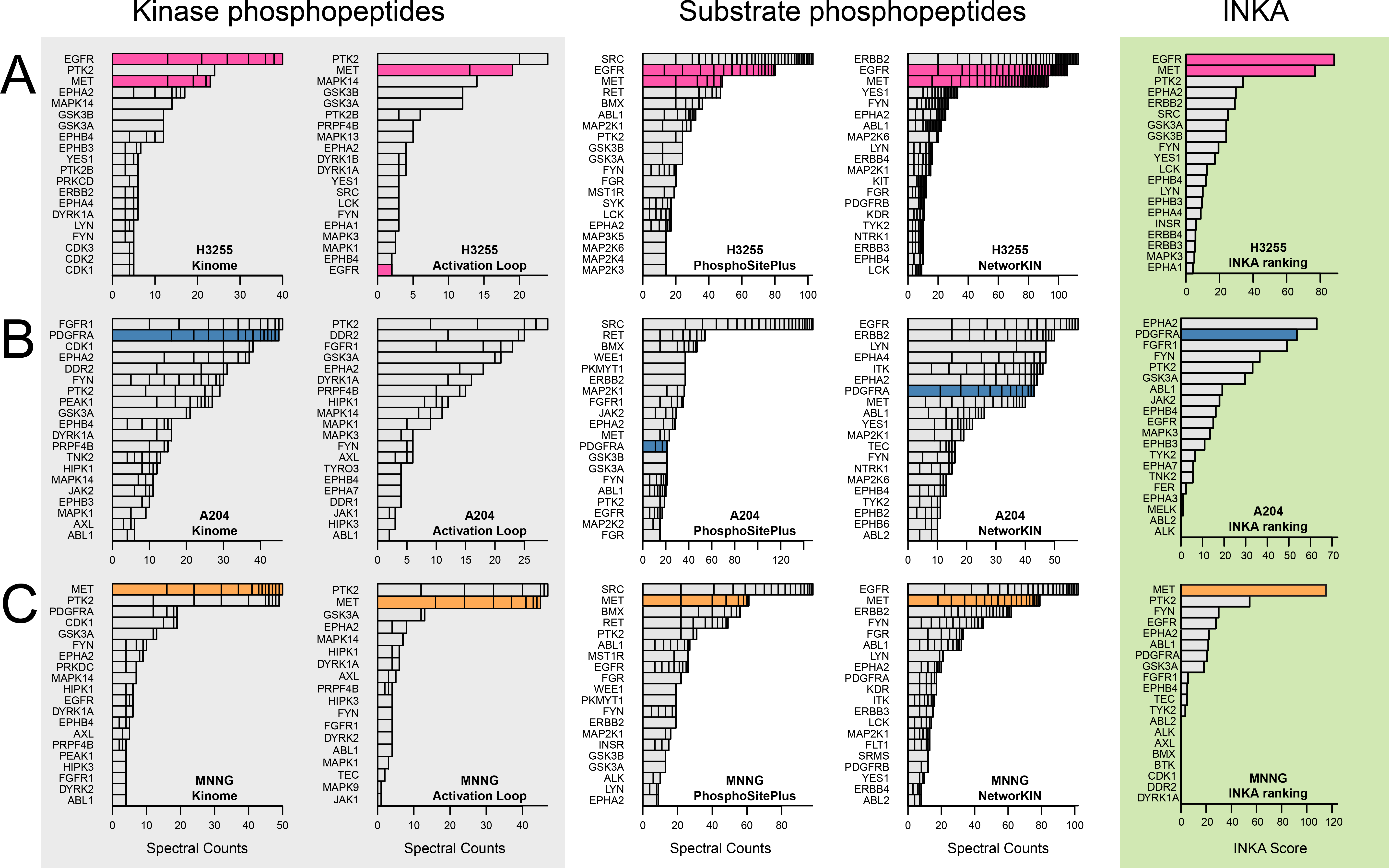
INKA analysis of three cancer cell line cases from the literature. A H3255 NSCLC cells with an *EGFR* mutation. INKA analysis shows that EGFR is a hyper-activated kinase (top 2 in all branches), together with MET. B A204 rhabdomyosarcoma cells with documented PDGFRA signaling. PDGFRA exhibits variable ranking (top-intermediate) in individual analysis types for A204, but integrated INKA analysis (right-most bar graph) infers it as a highly active, rank-2 kinase, after EPHA1. C MNNG/HOS osteosarcoma cells with documented MET signaling. MET consistently ranks among the top 2 in all analysis arms, culminating in a first rank in the integrative INKA analysis. Data information: See the legend of Figure 2 for basic explanation.

Altogether, analysis of public datasets illustrates the capacity of INKA scoring to identify kinases that are relevant oncogenic drivers in diverse cancer cell lines at baseline.

### Testing the INKA approach in differential settings

To explore the discriminative power of INKA scoring, we analyzed pTyr-phosphoproteomic data from wild-type versus mutant cells, and from untreated versus drug-treated cells as genetic and pharmacological dichotomies (Fig 5, Supplementary Table 3). First, we utilized raw data from our laboratory (van der Mijn *et al*, 2014) to compare wild-type U87 glioblastoma cells with isogenic U87-EGFRvIII cells overexpressing a constitutively active EGFR mutant. The EGFR INKA score was significantly higher and dominating in U87-EGFRvIII relative to U87 (Fig 5A). Some other kinases also exhibited higher-ranking INKA scores, including MET and EPHA2, for which enhanced phosphorylation in U87-EGFRvIII was previously documented (Huang *et al,* 2007; Stommel *et al*, 2007), as well as SRC-family members. In a treatment setting, INKA analysis clearly revealed a specific drug effect after targeting EGFR in U87-EGFRvIII, with the high, first-rank INKA score for EGFR at baseline being halved after treatment with erlotinib (Fig 5B).

**Figure 5.**
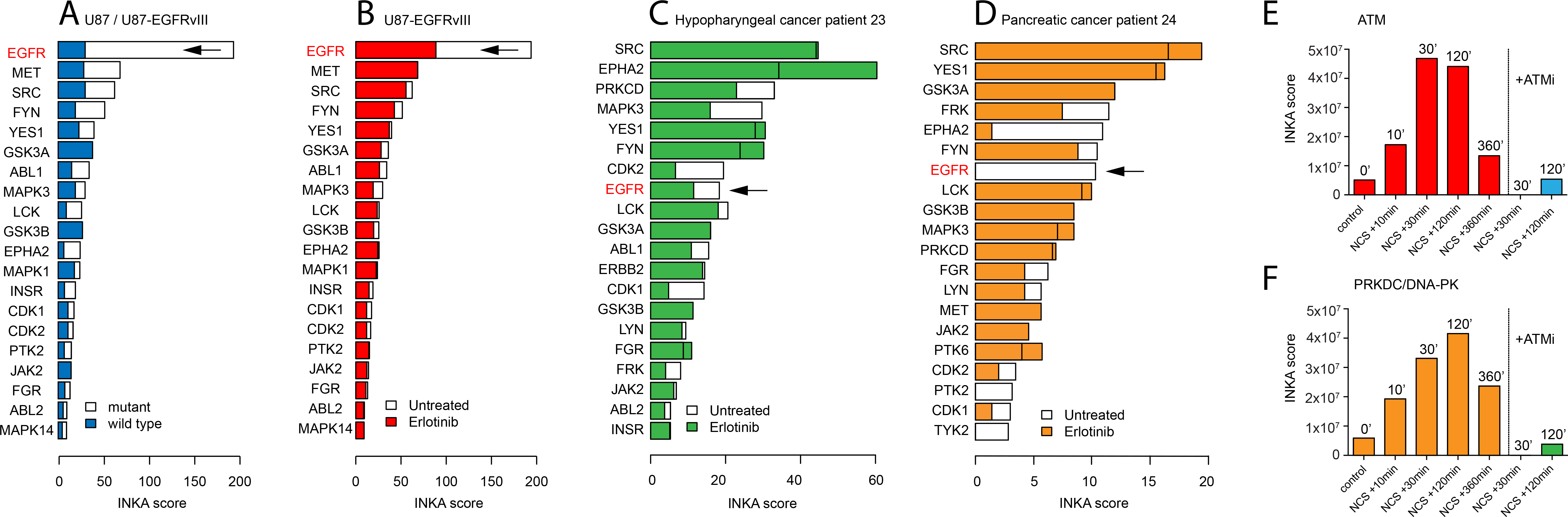
INKA analysis in differential genetic and pharmacological settings. A Effect of a monogenetic change in a cancer cell line use case. Comparison of U87 glioblastoma cells (“wild type”) with isogenic U87-EGFRvIII cells overexpressing a consitutively active EGFR variant (“mutant”) grown under baseline conditions. B Effect of drug treatment in a cancer cell line use case. Comparison of U87-EGFRvIII cells at baseline with U87-EGFRvIII cells treated with 10 µM erlotinib for 2 h shows a clearly reduced INKA score for EGFR. C Effect of drug treatment in a patient with hypopharyngeal cancer. Tumor biopsies were taken both before and after two weeks of erlotinib treatment. D Same as panel C, but for a patient with pancreatic cancer. E Time-dependent effect of radiomimetic treatment in a cancer cell line use case. MS intensity-based INKA analysis of TiO2-captured phosphoproteomes from G361 melanoma cells at different time points following treatment with the DNA damage-inducing drug neocarzinostatin (NCS) in the absence or presence of ATM inhibitor KU55933 (ATM). Plotted is the INKA score for ATM, exhibiting a time-dependent increase, which is not observed with ATM blocking. Full INKA score bar graphs are shown in Supplementary Fig **8**. F Same as panel E, but plotting of the INKA score for PRKDC/DNA-PK, which exhibits similar behavior. Raw data for panels A&B are from Van der Mijn *et al.* (van der Mijn *et al*, 2014). Raw data for panels E&F are from Bensimon *et al.* (Bensimon *et al*, 2010) and averaged for replicate treatment conditions.

Second, in order to extend these findings to a clinical framework, INKA scoring was applied to pTyr-phosphoproteomic data on patient tumor biopsies (Fig 5C,D). Biopsies were collected both before and after two weeks of erlotinib treatment (standard dose, clinical trial NCT01636908; Labots *et al.*, submitted for publication). Importantly, the on-treatment biopsy from a patient with advanced head and neck squamous cell carcinoma showed a reduced INKA score and rank for EGFR as well as cell cycle-associated kinases (Fig 5C). Interestingly, in a pancreatic cancer patient, no residual EGFR activity could be inferred by INKA in a tumor biopsy after erlotinib treatment (Fig 5D).

Third, we assessed performance of INKA analysis using published phosphoproteome data (Bensimon *et al*, 2010) of human G361 melanoma cells after induction of genotoxic stress with neocarzinostatin, a radiomimetic that induces double-strand breaks (Fig 5E,F). As these data were generated using an earlier-generation mass spectrometer that is less sensitive than current orbitrap-based instruments, count data did not work well, and MS intensity data were analyzed instead. As expected in the context of DNA damage signaling, INKA scores for ATM and PRKDC/DNA-PK exhibited a time-dependent increase after addition of neocarzinostatin (Fig 5E,F and Supplementary Fig 8). Moreover, the ATM INKA score was significantly reduced after the addition of ATM inhibitor KU55933 (Fig 5E). INKA scoring suggests that the inhibitor influences PRKDC as well (Fig 5F).

In summary, the above differential analyses of phosphoproteomes show that the INKA pipeline can pinpoint target activation and inhibition after perturbation in both cell lines and clinical samples.

## Discussion

Present-day cancer treatment is increasingly shifting towards individualized therapy by specific targeting of hyperactive kinases in patient tumors. In this context, with heterogeneity and plasticity of kinase signaling in a specific tumor at a specific time, it is essential to have an overview of (hyper)active kinases and prioritize ones that (help) drive malignancy to maximize therapeutic success and minimize expensive failures and unnecessary burden for the patient. Here, we present a novel pipeline, Integrative Inferred Kinase Activity (INKA) scoring, to investigate phosphoproteomic data from a single sample and identify (hyper)active kinases as candidates for (co-)targeting with kinase inhibitors. In a first demonstration of its application, we have performed INKA analyses of established cancer cell lines with known oncogenic drivers. We analyzed both data from tyrosine phosphoproteomics in our laboratory and similar data described in the literature (five cell lines in each case). Furthermore, INKA could distinguish relevant differences between closely related mutant and wild type cells and reveal drug perturbation effects in both tyrosine and global (TiO_2_) phosphoproteomics data. We also applied INKA scoring to tumor needle biopsies of two patients before and after kinase inhibitor treatment.

This study shows that data from label-free MS-based phosphoproteomics can be harnessed in a multipronged analysis to infer and rank kinase activity in individual biological samples. In kinase-centric analyses, phosphorylation of the kinase itself (either considering all sites or focusing on the ones located in the activation loop) is used as a proxy for its activation. In substrate-centric analyses, instead, substrate phosphorylation is used to deduce kinase activity indirectly through kinase-substrate relationships (based on either experimental knowledge or motif-based prediction). Here we developed INKA that combines kinase-centric and substrate-centric evidence in a stringent meta-analysis, to yield an integrated metric for inferred kinase activity. The results of INKA highlight kinases that are in line with known cancer biology and show that INKA scoring clearly outperforms substrate-centric analyses alone, and also holds a slight edge over phosphokinase ranking pioneered by Rikova et al^12^. In particular, kinases driving tumor growth rank high in INKA scoring, illustrating the power of applying an integrative analysis of in-depth phosphoproteomics data.

Meaningful substrate-centric inference of kinase activity is pivotal to INKA scoring. It depends on the availability of comprehensive curated data on experimentally observed kinase-substrate relationships or reliable predictions thereof. To date, this requirement has only been partly fulfilled, and more substantial inferences could be made with further population of resources (such as PhosphoSitePlus), especially when the latter also cover cancer-associated aberrations such as fusion genes BCR-ABL and EML4-ALK (Medves & Demoulin, 2012; Lee *et al*, 2017). To overcome current limitations in information, our INKA pipeline provides a separate kinase-centric ranking of kinases that are not yet covered by PhosphoSitePlus and NetworKIN, but do show up as phosphoproteins in a sample. This reduces the chance of missing important kinases, as illustrated by the case of AXL in the HCC827-ER3 cell line. Moreover, the INKA pipeline generates network visualizations of all kinase-substrate relationships inferred for the top 20 kinases in an experiment. This provides a more instructive overview of phosphoproteomic biology in a sample than mere scoring and ranking alone.

A next stage is to apply INKA to more advanced cancer models and, especially, clinical samples. To analyze limited amounts of patient tumor tissue in a clinical practice setting, we have recently downscaled tyrosine phosphoproteomics to clinical needle-biopsy levels^56^. Using this workflow, we performed proof-of-concept analysis of patient samples from a clinical molecular profiling study with kinase inhibitors (Labots et al., submitted), showing relevant effects.

In summary, INKA scoring can infer and rank kinase activity in a single biological sample, and display differences between closely related yet genetically distinct cells and after drug intervention. INKA analysis of phosphoproteomic data on tumor biopsies collected in kinase inhibitor trials can pave the way for future clinical application. The ultimate goal would be tailoring treatment selection for the individual patient.

## Materials and Methods

### Cell lines and culture

H2228 non-small cell lung cancer cells carry an *EML4-ALK* fusion gene, SK-Mel-28 melanoma cells harbor a V600E *BRAF* mutation, and K562 chronic myelogenous leukemia cells carry a *BCR-ABL1* fusion gene (philadelphia chromosome). HCC827 and HCC827-ER3 are non-small cell lung cancer cell lines with a deletion of EGFR residues E746-A750, the latter cell line being a subclone of HCC827 after selection for acquired resistance to erlotinib in a murine xenograft setting (Zhang *et al*, 2012). HCC827, H2228, SK-Mel-28 and K562 were obtained from the American Type Culture Collection (ATCC, Rockville, MD) and HCC827-ER3 was provided by Prof. Balazs Halmos, Division of Hematology/Oncology, Columbia University Medical Center, New York, USA.

H2228 and SK-Mel-28 were cultured in 175-cm^2^ flasks containing DMEM supplemented with 10% fetal bovine serum and 2mM L-glutamine. K562 was cultured as a suspension in RPMI 1640 containing 2 mM L-glutamine and supplemented with 10% fetal bovine serum. HCC827 and HCC827-ER3 were grown in 15-cm dishes with the latter medium. Cell lines tested negative for mycoplasma.

### Cell lysis and phosphoproteomics

Cells were grown to 70-80% confluency (adherent cell lines) or to a 1 x 10^6^ cells/ml suspension (K562), washed with phosphate-buffered saline (PBS), lysed at approximately 2˓ x 10^7^ cells/ml in 5 ml lysis buffer (9 M urea, 20 mM HEPES pH 8.0, 1 mM sodium orthovanadate, 2.5 mM sodium pyrophosphate, 1mM ß-glycerophosphate), sonicated (3 cycles of 30 s), and extracts were stored at −80 °C.

For phosphoproteomics, lysate aliquots equivalent to 10 mg total protein were used as described before (van der Mijn *et al*, 2015). Proteins were reduced by incubation in 4.5 mM dithiothreitol for 30 min at 55 °C, alkylated in 10 mM iodoacetamide for 15 min at room temperature in the dark, and digested overnight at room temperature with 10 µg/ml trypsin in a fourfold increased volume containing 20 mM HEPES pH 8.0. After acidification (trifluoroacetic acid to 1% final concentration), tryptic digests were desalted on Sep-Pak C18 cartridges (Waters Chromatography, Etten-Leur, Netherlands) and lyophilized. Peptides were taken up in 700 µl immunoprecipitation buffer (50 mM MOPS pH 7.2, 10 mM sodium phosphate, 50 mM NaCl) and transferred at 4 °C to a microcentrifuge tube containing 40 µl of a 50% (v/v) slurry of agarose beads harboring P-Tyr-1000 anti-phosphotyrosine monoclonal antibodies (Cell Signaling Technologies, Danvers, USA) that had been washed and taken up in PBS. Following a 2-h incubation at 4 °C on a rotator, beads were washed twice with cold PBS, and three times with cold HPLC-grade water. Bound peptides were eluted with a total of 50 µl 0.15% trifluoroacetic acid in two steps. After desalting with custom-made C18 stage tips, drying, and solubilization in 20 µl 4% acetonitrile/0.5% trifluoroacetic acid, samples were subjected to LC-MS/MS analysis. For tumor biopsy phosphoproteomics, tumor needle biopsies from a patient with advanced head and neck squamous cell carcinoma, obtained in an Institutional Review Board-approved molecular profiling study before and after 2 weeks of treatment with erlotinib (NCT clinical trials identifier 01636908; www.clinicaltrials.gov), were cut and processed as described elsewehere (Labots *et al*, 2017). Peptide preparation from tumor biopsy lysates was performed with 2.5 mg protein input for both pre- and on-treatment biopsies.

### LC-MS/MS

Peptides were separated on an Ultimate 3000 nanoLC-MS/MS system (Dionex LC-Packings, Amsterdam, The Netherlands) equipped with a 20-cm, 75-µm inner diameter fused silica column, custom packed with 1.9-µm ReproSil-Pur C18-AQ silica beads (120-Å pore size; Dr. Maisch, Ammerbuch-Entringen, Germany). After injection, peptides were trapped at 6 µl/min on a 10-mm, 100-µm inner diameter trap column packed with 5-µm ReproSil-Pur C18-AQ silica beads (120-Å pore size) in buffer A (buffer A: 0.5% acetic acid, buffer B: 80% acetonitrile, 0.5% acetic acid), and separated at 300 nl/min with a 10–40% buffer B gradient in 90 min (120 min inject-to-inject). Eluting peptides were ionized at a potential of +2 kV and introduced into a Q Exactive mass spectrometer (Thermo Fisher, Bremen, Germany). Intact masses were measured in the orbitrap with a resolution of 70,000 (at m/z 200) using an automatic gain control (AGC) target value of 3 × 10^6^ charges. Peptides with the top 10 highest signals (charge states 2+ and higher) were submitted to MS/MS in the higher-energy collision cell (4-Da isolation width, 25% normalized collision energy). MS/MS spectra were acquired in the orbitrap with a resolution of 17,500 (at m/z 200) using an AGC target value of 2 × 10^5^ charges and an underfill ratio of 0.1%. Dynamic exclusion was applied with a repeat count of 1 and an exclusion time of 30 s.

### Peptide identification and quantification

MS/MS spectra were searched against theoretical spectra based on a UniProt complete human proteome FASTA file (release January 2014, no fragments; 42104 entries) using MaxQuant 1.4.1.2 software (Cox & Mann, 2008). Enzyme specificity was set to trypsin, and up to two missed cleavages were allowed. Cysteine carboxamidomethylation (+57.021464 Da) was treated as fixed modification and serine, threonine and tyrosine phosphorylation (+79.966330 Da), methionine oxidation (+15.994915 Da) and N-terminal acetylation (+42.010565 Da) as variable modifications. Peptide precursor ions were searched with a maximum mass deviation of 4.5 ppm, and fragment ions with a maximum mass deviation of 20 ppm. Peptide and protein identifications were filtered at a false discovery rate of 1% using a decoy database strategy. The minimal peptide length was set at 7 amino acids, the minimum Andromeda score for modified peptides was 40, and the corresponding minimum delta score was 17. Proteins that could not be differentiated based on MS/MS spectra alone were clustered into protein groups (default MaxQuant settings). Peptide identifications were propagated across samples using the ‘match between runs’ option. Phosphopeptide MS/MS spectral counts (Liu *et al*, 2004) were calculated from the MaxQuant evidence file using R.

### INKA analysis

The INKA analysis pipeline was implemented in R, utilizing custom tables with data extracted from web resources including UniProt (UniProt Consortium, 2015) (for mapping attributes of UniProt accessions; www.uniprot.org, mapping date 8 June 2016), PhosphoSitePlus (Hornbeck *et al*, 2015) (for experimentally observed phosphorylation sites and kinase-substrate relationships; www.phosphosite.org, Phosphorylation_site_dataset and Kinase_Substrate_Dataset, versions of 3 July 2016), KinBase (Manning *et al*, 2002) (for currently recognized protein kinases; kinase.com/web/current/kinbase, mapping date 20 July 2016), and HGNC (Yates *et al*, 2017) (for mapping to official gene symbols of the HUGO Gene Nomenclature Committee; www.genenames.org). Furthermore, a tool for the identification of kinase activation loop peptides (Munk *et al*, 2016), provided on the Phomics website (phomics.jensenlab.org/ activation loop peptides), and a locally running version of the NetworKIN algorithm (Horn *et al*, 2014) were used. For the latter NetworKIN engine, NetworKIN version 3.0 code was used in combination with a UniProt human reference proteome FASTA file derived from release 2014_01 filtered for “no fragments”, and containing 21849 TrEMBL entries and 39703 Swiss-Prot entries.

#### Data filtering and annotation

For phosphopeptide data, we used data from the MaxQuant ‘modificationSpecificPeptides’ table, combined with spectral count data calculated above. Although INKA analysis can be performed with intensity-based quantification, we favor spectral count-based quantification as it is less sensitive to peptides with outlier intensities and is more robust for the analysis of aggregated data for multiple peptides, some of which may exhibit dominantly high intensities. For our QExactive data, spectral counting outperformed intensity-based quantification for INKA-based kinase ranking of known drivers (data not shown), yet for the low level LTQ-FTMS data the intensity data worked better (Fig 5E,F; count data not shown). Table rows with data linked to multiple UniProt gene symbols were deconvoluted into separate rows with a single gene symbol. For phosphosite data, we used data from the MaxQuant ‘Phospho (STY)Sites’ table, filtering for so-called class I sites (localization probability > 0.75). Table rows with data linked to multiple UniProt accessions, and those linked to multiple phosphopeptides, were deconvoluted into separate rows. Data from the web resources mentioned above were used to prioritize rows linking the same phosphosite to the same gene, only retaining the row with the best annotated accession. Subsequently, the phosphosite data were merged with pertinent phosphopeptide data in a single, non-redundant class-I phosphosite-phosphopeptide table.

#### Plot data generation

(1) Kinome analysis: A table with data for kinome analysis was generated from the deconvoluted phosphopeptide table obtained above by removing rows with different UniProt gene symbols linking the same phosphopeptide to the same HGNC-mapped symbol, and then filtering for rows with phosphopeptides derived from protein kinases (HGNC-mapped symbol among those for established protein kinases catalogued in KinBase). (2) Activation Loop analysis: the table generated for kinome analysis was further filtered for phosphopeptides that harbor activation loop sites as indicated by the Phomics web resource. (3) PhosphoSitePlus (PSP) analysis: the non-redundant class-I phosphosite-phosphopeptide table was merged with a table with data from PhosphoSitePlus detailing experimentally observed human kinase-substrate relationships. (4) NetworKIN (NWK) analysis: based on phosphosite data in the non-redundant class-I phosphosite-phosphopeptide table, kinase-substrate relationships were predicted (outside R) using the NetworKIN algorithm in combination with the same FASTA file that was used for peptide and protein identification. The results were imported in our R pipeline and restricted to predictions associated with a NetworKIN score of at least 2 that, in addition, exceeded 90% of the score of the top prediction for the same phosphosite. The non-redundant class-I phosphosite-phosphopeptide table was then merged with the filtered NetworKIN results table.

From the four analysis tables, lists with plot data tables for individual samples were generated as follows. For the kinase-centric kinome and activation loop analyses, per sample, kinase gene symbols and associated spectral count values were extracted from all rows (corresponding to phosphopeptides) in the respective tables, and spectral counts were multiplied by the number of phosphorylations of the pertinent peptide so as to account linearly for all phosphorylation activity impinging on the protein. For the substrate-centric PSP and NWK analyses, the same was done with the corresponding analysis tables, but multiplied spectral counts were further divided by the number of phosphosites inferred by MaxQuant in the ‘Phospho (STY)Sites’ table. As the latter number may exceed the number of actual phosphomodifications of the phosphopeptide, this makes sure the signal is not exaggerated.

#### Kinase bar graph plotting

To visualize kinase phosphorylation levels (kinase-centric analyses), or substrate phosphorylation levels attributed to specific kinases (substrate-centric analyses), the above plot data lists were used to produce, per sample, bar graphs for the top 20 kinases in each of the four analyses. Specifically, all signals (spectral counts) assigned to a specific kinase were represented by stacked bar segments. For kinase-centric analyses, bar segments correspond to individual phosphopeptides; for substrate-centric analyses, they represent individual phosphosites (aggregated signal of one or more phosphopeptides).

#### INKA analysis and plotting

To infer kinase activity within a single sample, we integrated data from the four analysis tables relevant to the sample (visualized separately in kinase bar graphs above), and calculated, for each kinase, a single INKA score metric. First, from the different analyses we aggregated cumulative signals (corresponding to bars above) assigned to a given kinase, summing values from the kinome analysis and those from the activation loop analysis to get a kinome-centric measure (*C_kin_*) according to equation (1), and summing values from the PSP and NWK analyses to get a substrate-centric measure (*C_sub_*) according to equation (2):
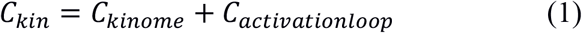

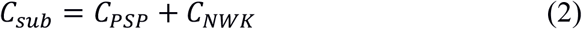

Subsequently, the geometric mean of these two measures was taken to obtain the final, integrated INKA score via equation (3), and a ’skew’ parameter was defined in equation (4) to indicate the relative contribution of kinase-centric versus substrate-centric evidence to the INKA score:
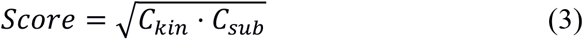

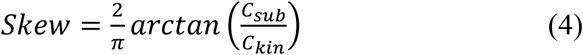

The skew parameter is a goniometric correlate of the ratio of substrate-centric and kinase-centric evidence, with the corresponding angle normalized on 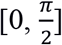 to give a value in [0,1]. The value is 0 when all evidence is kinase-centric, 1 when all evidence is substrate-centric, and 0.5 when there is equal contribution from both sides. To produce an intuitive, visual representation of the inferred (hyper)active kinases in a given experimental sample, for each kinase, the combination of the INKA score and skew parameter values were taken as vertical and horizontal coordinates, respectively, to create a scatter plot.

Kinases for which there is no available data on experimentally observed substrates (PSP), nor any reliable sequence motif with which to predict the kinase’s substrates (NWK), will always get a zero *C_sub_* score, and therefore a zero INKA score. Therefore, for these kinases we performed a separate ranking solely based on the kinase-centric side of the INKA score, i.e, the sum of signals in the kinome and activation loop analyses, *C_kin_*. Flanking the INKA scatter plot, a bar graph was plotted for the top 20 of such ‘out-of-scope’ kinases, only considering kinases associated with a total of at least 2 spectral counts. As a more simplified visualization, INKA scores for the top 20 kinases were also plotted in a bar graph for each sample.

#### Kinase-substrate relationship network plotting

For network visualization of inferred kinase-substrate relations annotated with INKA scores, the ‘network’ R package (Butts, 2008) was used. Per sample, relevant data were extracted from a merge of the PSP and NWK analysis tables generated above. Edge lists were created with kinases and substrates as tail and head nodes, respectively, of directed edges. Other data were stored as node and edge attributes in the network object. Nodes were depicted as a hexagon for observed kinases (one or more phosphopeptides derived from the kinase identified), as a pentagon for inferred kinases (no direct observation, but linked to phosphorylation of one or more substrate phosphopeptides), or as a circle for non-kinase substrates. INKA scores were used to define kinase node colors in a white to red gradient, and kinases with at least one phosphorylated activation loop phosphosite were indicated by a thicker node border. Edge widths were made to correlate with the associated substrate site phosphorylation level, and edge colors to the analysis on which the kinase-substrate relationship was based (PSP: coral, NWK: blue, both: green). Node coordinates were calculated using a Fruchterman-Reingold layout algorithm for force-directed graph drawing with empirically chosen parameter settings (niter=100N, area=N^1.8^, repulse.rad=N^1.5^, ncell=N^3^; N: number of nodes).

#### Statistical significance assessment

Random INKA scores for each kinase in each sample were generated using a two-fold randomization procedure. First, all measured spectral count values were permuted in a sample-wise fashion. Second, as the kinase responsible for phosphorylation of a specific phosphosite, a kinase was randomly selected from the pool of all kinases that could possibly be inferred. To this end, all kinase-substrate links in the PhosphoSitePlus data table and in a proteome-wide NetworKIN analysis table (obtained for all phosphosites in all protein sequences contained in the same FASTA file as used in the identification stages) were randomized. The result of this procedure was then used as input for the calculation of INKA scores in the same way as described above. Iterating this procedure 100,000 times gave sample-and kinase-specific null distributions which enabled the calculation of p-values for each of the original INKA scores associated with each sample.

## Acknowledgements

SK-Mel-28 cells from ATCC were kindly provided by Dr. Elisa Giovannetti, AIRC/Start-Up Unit, Pisa University, Italy. HCC827 ER3 cells were kindly provided by Prof. Balazs Halmos, Division of Hematology/Oncology, Columbia University Medical Center, New York, USA. Raw label-free phosphoproteomic data on human melanoma G361 cells after radiomimetic treatment were kindly provided by Dr. A. Bensimon, ETH Zürich, Switzerland. Inge de Reus is acknowledged for experimental assistance. Dr. Frank Koopman is thanked for critically reading the manuscript. VitrOmics Healthcare Services (VHS), Cancer Center Amsterdam, and NWO-Middelgroot (project number 91116017) are acknowledged for support of the mass spectrometry infrastructure, and VHS also for support of RB.

## Author Contributions

CRJ designed the experiments. TVP, RB, JCK, AH wrote the R script and performed the data analysis. AH proposed the combination score and developed the web server. CvA and RdH performed the experiments. CRJ, CvA and TLL gave input on the analysis. EH and SRP optimized NetworKIN settings. SRP performed nanoLC-MS/MS, database searching and prepared figures. HV and ML provided clinical expertise. CRJ, SRP, TVP, RB, CvA, JCK, AH, FR, ML and HV wrote the paper.

## Data Availability

The mass spectrometry data from this publication have been deposited to the ProteomeXchange Consortium via the PRIDE partner repository [www.ebi.ac.uk/pride/archive] and assigned the identifier PXD006616.

## Code Availability

Researchers can analyze their data by the INKA pipeline via www.inkascore.org.

## Supplementary Figure Legends

**Supplementary Figure 1. Coverage of 538 unique protein kinases from KinBase by resources providing kinase-substrate relationships.**

The Venn diagram shows that 172 kinases in KinBase are missing from both PhosphoSitePlus (PSP) and NetworKIN (NWK), 31 are only covered by NWK, 172 are only covered by PSP, while 163 are covered by both NWK and PSP. Additionally, 14 proteins annotated by PSP are not present in KinBase, but mostly involve small-molecule kinases or proteins with spurious (electronic) annotation, with the exception of two fusion protein species with established protein kinase activity (BCR-ABL and NPM-ALK).

**Supplementary Figure 2. Correlation between kinase-specific INKA scores and estimated p-values.**

Log10-transformed INKA scores are plotted against (sign-switched) log10-transformed p-values for each of four cell line use cases. Highly significant, positive Spearman rank correlation coefficients for each of the cell lines demonstrate a monotonic trend: higher INKA scores are associated with lower p-values. (a) K562 chronic myelogenous leukemia cells. (b) SK-Mel-28 melanoma cells. (c) HCC827-ER3 lung cancer cells. (d) H2228 lung cancer cells.

**Supplementary Figure 3. Kinase-substrate relation network for top 20 INKA-scoring kinases and their observed substrates in K-562 chronic myelogenous leukemia cells with a *BCR-ABL* fusion.**

ABL1 is a highly connected and central node. Kinases downstream of BCR-ABL signaling, such as SRC, are also active, albeit to a lower extent.

**Supplementary Figure 4. Kinase-substrate relation network for top 20 INKA-scoring kinases and their observed substrates in SK-Mel-28 melanoma cells with BRAF^V600E^.**

Two clusters of activated kinases are observed, one containing BRAF targets MAPK1 and MAPK3, and the other containing SRC as highly connected nodes.

**Supplementary Figure 5. Kinase-substrate relation network for top 20 INKA-scoring kinases and their observed substrates in HCC827-ER3 non-small cell lung carcinoma cells.**

EGFR and MET are central and highly connected nodes. AXL, associated with erlotinib resistance of HCC827-ER3 cells, is missed by INKA score-based analysis as no substrate-centric data is available for this kinase.

**Supplementary Figure 6. INKA plot and top 20 INKA-scoring kinase bar graph for HCC827 cells with mutant EGFR.**

In the INKA plot proper, the vertical position of kinases (driver in red) is determined by their INKA score, whereas the horizontal position is determined by the (im)balance of evidence from kinase-centric and substrate-inferred arms of the analysis. EGFR and MET are implicated as the most active kinases. Kinases not covered by PhosphoSitePlus (PSP) and NetworKIN (NWK) are visualized in a flanking bar graph on the left. Note absence from the bar graph of the AXL kinase that is responsible for erlotinib resistance in the HCC827-ER3 sub-line (see Fig 3c).

**Supplementary Figure 7. Kinase-substrate relation network for top 20 INKA-scoring kinases and their observed substrates in H2228 non-small cell lung carcinoma cells with an *EML4-ALK* fusion.**

Multiple highly active and connected nodes are present in the network for H2228, implying relative insensitivity to inhibition of ALK alone, in line with previous functional data. Dual inhibition of the number-5 hyperactive node, EGFR, and ALK results in significant reduction of proliferation (Voena *et al*, 2013).

**Supplementary Figure 8. MS intensity-based INKA analysis of data published by Bensimon et al. on TiO_2_-captured phosphoproteomes from G361 melanoma cells following radiomimetic treatment.**

A INKA score bar graphs for G361 at baseline.

B-E INKA score bar graphs for G361 after 10 min (B), 30 min (C), 120 min (D), or 360 min (E) treatment with 200 ng/ml neocarzinostatin (NCS).

F,G INKA score bar graphs for G361 after 30 min (F) or 120 min (G) treatment with 200 ng/ml neocarzinostatin in the presence of 10 µM KU55933 (ATM inhibitor, ATMi). DNA damage-induced kinases ATM, ATR, and PRKDC/DNA-PK are highlighted in red.

Data information: raw data (Bensimon *et al*, 2010) were normalized and averaged for replicate treatment conditions.

## Supplementary Table Legends

**Supplementary Table 1. Mass-spectrometric and annotation data for phosphopeptides that were captured and identified in phosphotyrosine-based immunoprecipitation experiments on cell lines and patient tumors.**

Pair-wise tables “phosphopeptides…” and “phosphosites…” harbor information from selected columns of the MaxQuant modificationSpecificPeptides.txt and Phospho (STY)Sites.txt export files, respectively. The latter list phosphosites inferred by MaxQuant on the basis of mass-spectrometric evidence. Raw intensities were median normalized, and MS/MS spectral counts were obtained with the help of the MaxQuant evidence.txt export file.

**Supplementary Table 2. Coverage of protein kinases by various resources used in this study.**

A 538 established protein kinases from KinBase (http://kinase.com/web/current/kinbase)

B 489 kinases for which activation loop segment annotation is available from Phomics (http://phomics.jensenlab.org)

C 349 kinases linked to substrate phosphorylation in experimental studies, provided by PhosphoSitePlus (http://www.phosphosite.org/staticDownloads.action)

D 194 kinases linked to substrate phosphorylation through prediction, provided by NetworKIN (http://networkin.info)

E 172 protein kinases that are not covered by both PhosphoSitePlus and NetworKIN F 14 entries from PhosphoSitePlus that are not covered by KinBase.

**Supplementary Table 3. Quantitative result tables for INKA analyses.**

A-O Detailed are analyses for (A) K562 chronic myeloid leukemia cells, (B) SK-Mel-28 melanoma cells, (C) HCC827-ER3 non-small cell lung carcinoma cells; sub-line of HCC827, (D) H2228 non-small cell lung carcinoma cells. (E) HCC827 non-small cell lung carcinoma cells; parental line of HCC827-ER3, (F) H3255 non-small cell lung carcinoma cells (Guo *et al*, 2008), (G) A204 rhabdomyosarcoma cells (Bai *et al*, 2012), (H) MNNG/HOS osteosarcoma cells (Bai *et al*, 2012), (I) U87 glioblastoma cells (van der Mijn *et al*, 2014), (J) U87-EGFRvIII glioblastoma cells (van der Mijn *et al*, 2014), (K) U87-EGFRvIII glioblastoma cells treated with erlotinib (van der Mijn *et al*, 2014), (L) tumor tissue biopsy from a hypopharyngeal cancer patient (Pt23) before treatment, (M) tumor tissue biopsy from a hypopharyngeal cancer patient (Pt23) after two weeks of erlotinib treatment, (N) tumor tissue biopsy from a pancreatic cancer patient (Pt24) before treatment, (O) tumor tissue biopsy from a pancreatic cancer patient (Pt24) after two weeks of erlotinib treatment.

Data information: For each given kinase, metrics are given for individual analyses (Kinome, ActLoop, PSP, and NWK), accompanied by kinase-centric and substrate-centric INKA score components (Kin = Kinome + ActLoop, Sub = PSP + NWK), INKA scores (Score = √ (Kin * Sub)), relative INKA scores (Rel.Score) and skew parameters indicating relative contribution from kinase-centric versus substrate-centric evidence (Skew = arctan (Sub/Kin) * 2/*π*). Skew ranges from 0 (kinase-centric evidence only) to 1 (substrate-centric evidence only), so that a skew of 0.5 indicates equal contribution.

**Supplementary Table 4. Drug sensitivity data for top 5 INKA-scoring kinases in four cancer cell line use cases.**

A K562 driven by BCR-ABL. The data show high sensitivity of the cell line only to inhibitors of ABL.

B SK-Mel-28 driven by BRAF. Drug inhibition data are also listed for BRAF and ERK-activating MEKs in addition to top INKA-scoring kinases. Among the latter, ERK1 and ERK2 exhibit limited sensitivity to inhibitors. Direct targeting of BRAF, or (upstream of ERK1/2) MEK1/2, is much more successful, in line with BRAF driver function.

C HCC827 driven by EGFR. HCC827, the erlotinib-sensitive parental line of the erlotinib-resistant HCC827-ER3 sub-line, exhibits high sensitivity to EGFR inhibitors, but not MET inhibitors.

D H2228 associated with EML4-ALK. The data show moderate sensitivity of H2228 to ALK inhibitors, but little efficacy of inhibitors targeting top INKA-scoring kinases.

Data information: data were derived from the Genomics of Drug Sensitivity in Cancer website (GDSC, http://www.cancerrxgene.org) and include cell line-specific IC50 values and Z-scores for relative sensitivity to a given drug compared to all other cell lines.

